# Temporal dynamics of proteome and phosphorproteome during neuronal differentiation in the reference KOLF2.1J iPSC line

**DOI:** 10.1101/2025.03.25.645331

**Authors:** Ying Hao, Ziyi Li, Erika Lara, Daniel M. Ramos, Marianist Santiana, Benjamin Jin, Jacob Epstein, Jasmin Camacho, Nicole Carmiol, Isabelle Kowal, Paige Jarreau, Cory A. Weller, Sydney Klaisner, Laurel A. Screven, Caroline B. Pantazis, Mike A Nalls, Priyanka Narayan, Luigi Ferrucci, Andrew B. Singleton, Michael E. Ward, Mark R. Cookson, Yue Andy Qi

**Author notes:** These authors contributed equally (Ying Hao, Ziyi Li). Corresponding author: Dr. Yue Andy Qi.

## Abstract

Induced pluripotent stem cell (iPSC)-derived neurons have emerged as a powerful model to investigate both neuronal development and neurodegenerative diseases. Although transcriptomics and imaging have been applied to characterize neuronal development signatures, comprehensive datasets of protein and post-translational modifications (PTMs) are not readily available. Here, we applied quantitative proteomics and phosphoproteomics to profile the differentiation of the KOLF2.1J iPSC line, the first reference line of the iPSC Neurodegenerative Disease Initiative (iNDI) project. We developed an automated workflow enabling high-coverage enrichment of proteins and phosphoproteins. Our results revealed molecular signatures across proteomic and phosphoproteomic landscapes during differentiation of iPSC-derived neurons. Proteomic data highlighted distinct changes in mitochondrial pathways throughout the course of differentiation, while phosphoproteomics revealed specific regulatory dynamics in GTPase signaling pathways and microtubule proteins. Additionally, phosphosite dynamics exhibited discordant trends compared to protein expression, particularly in processes related to axon functions and RNA transport. Furthermore, we mapped the kinase dynamic changes that are critical for neuronal development and maturation. We developed an interactive Web app (https://niacard.shinyapps.io/Phosphoproteome/) to visualize temporal landscape dynamics of protein and phosphosite expression. By establishing baselines of proteomic and phosphoproteomic profiles for neuronal differentiation, this dataset offers a valuable resource for future research into neuronal development and neurodegenerative diseases using this reference iPSC line.

**Highlights:**

- Temporal dynamics of proteome and phosphoproteome profiles in KOLF2.1J iPSC derived neurons.
- Phosphoproteomics highlights GTPase signaling and microtubule regulation in neuronal differentiation.
- Kinome mapping reveals a shift in kinase activity patterns from early to late differentiation.
- Shinyapp for visualizing the trajectory of protein and phosphosite expression during neuronal differentiation.

## Introduction

Human induced pluripotent stem cell (iPSC)-derived neurons have become a powerful tool for studying neurodevelopment, neuronal function and disease mechanisms^1–3^. Although differentiation protocols recapitulate some aspects of neuronal development^4,5^, it is known that there is substantial variation between iPSC lines and protocols^6^. To improve reproducibility, we have extensively characterized the KOLF2.1J reference human iPSC line as part of the iPSC Neurodegenerative Disease Initiative (iNDI) at the Center for Alzheimer’s and Related Dementias (CARD) at the National Institutes of Health (NIH)^7^. Currently, this reference iPSC line has been widely adopted and applied in the neuroscience community, offering a reliable and standardized system for advancing neurodevelopmental biology^8,9,10^ and neurodegenerative disease research^11,12^.

Over the past decades, large-scale genomics and transcriptomic studies have provided quantitative insights into gene expression trajectories during neuronal development^13–15^. Recently, single-cell and single-nucleus RNA sequencing (sc, snRNA-seq) have been utilized to map the temporal and spatial dynamics of brain development^16,17^, and have highlighted the heterogeneity of differentiated neuron populations. Phosphoproteomic studies of mouse brain tissues have provided insights into the phosphorylation-dependent mechanisms regulating neuronal activity and synaptic functions^18,19^. Studies of hyperphosphorylation-induced phase transitions have shed light on the impact of excessive phosphorylation of motor proteins in neuronal cells, offering potential insights into neurodegenerative disorders^20^. Additionally, recent studies have leveraged the quantitative phosphoproteomics to uncover dysregulated kinase networks in Alzheimer’s disease^21,22^. However, the dynamics of protein and phosphorylation expression during neuronal differentiation remain underexplored.

Here, we integrated multiple robotic extraction and enrichment platforms into a single phosphoproteomics pipeline based around 96 well plates requiring minimal manual operation. Using this automated platform, we profiled the proteome and phosphoproteome of KOLF2.1J iPSCs over 28 days of neuronal differentiation. Integrative analysis unveiled stage-specific signatures and regulatory networks associated with neuronal maturation. For easy access and navigation of our longitudinal proteomic profiling of KOLF2.1J iPSC-derived neurons, we developed an interactive Shinyapp for data browsing (https://niacard.shinyapps.io/Phosphoproteome/). Our results provided a comprehensive understanding of the molecular mechanisms involved in neuronal development and offered an additional framework for interpreting results from the iNDI project.

## Results

### Temporal proteome and phosphoproteome profiling of KOLF2.1J-derived neurons

To investigate the molecular dynamics of neuronal differentiation, we differentiated KOLF2.1J iPSCs into mature neurons with the doxycycline-inducible Neurogenin-2 overexpression protocol^23^, and we performed proteomic and phosphoproteomic profiling of KOLF2.1J-derived neurons across six time points (Days [D] 0, 4, 7, 14, 21, and 28) (Figure 1A). Proteins and enriched phosphorylated peptides were prepared using an automated workflow for total proteomics (Figure 1B) and phosphoproteomics (Figure 1C), followed by data-independent acquisition (DIA)-based mass spectrometry acquisition and functional pathway analysis. Daily live-cell imaging of neurite outgrowth revealed rapid neurite extension between D4 and D26, followed by a slight decline at D28 (Figure 1D). This pattern suggests the presence of fully differentiated neurons at D25-D28, consistent with previous observations^7^.

**Figure 1.**
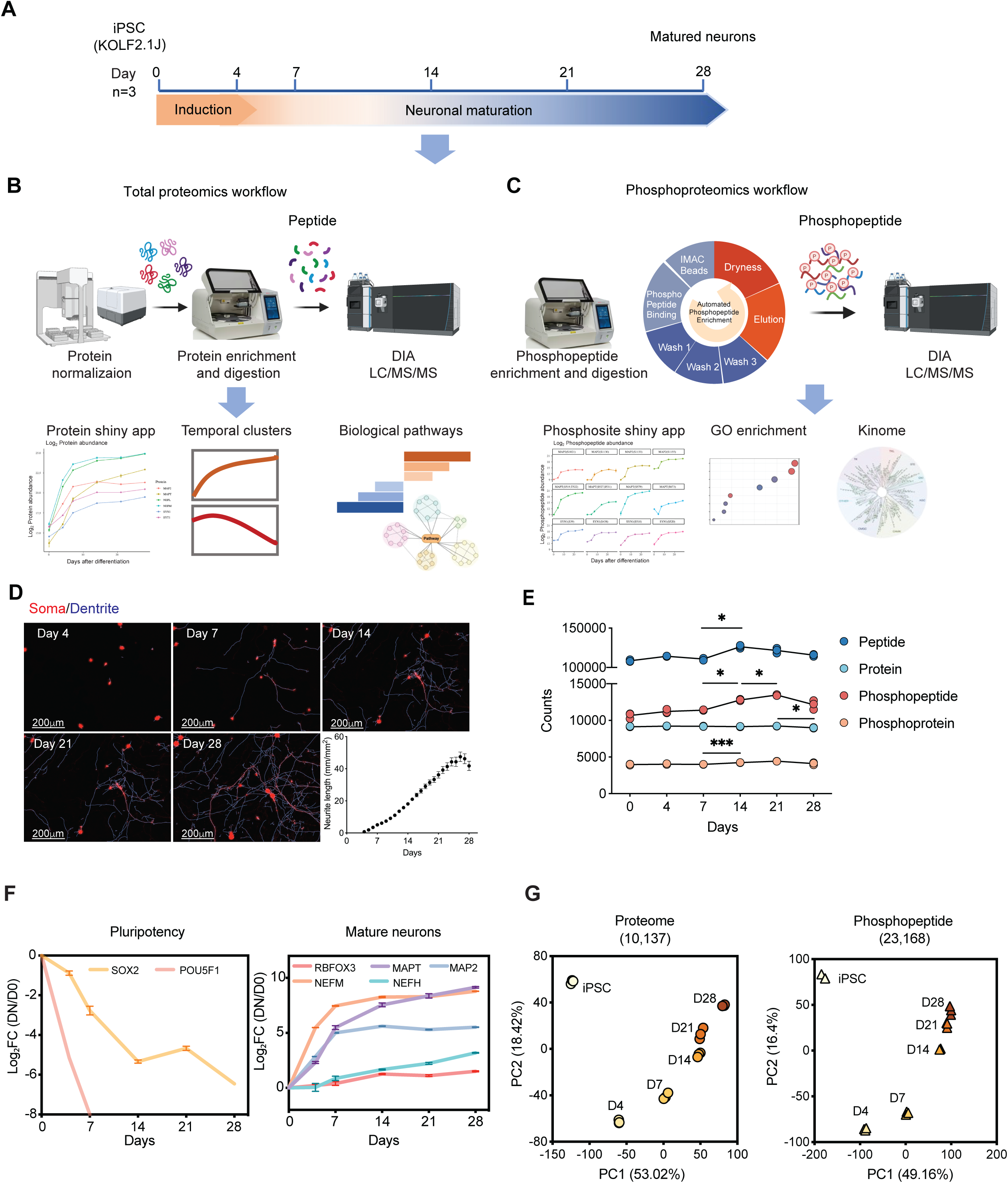
Temporal proteome and phosphoproteome profiling of KOLF2.1J-derived neurons. (A) Schematic overview of the workflow for neuronal differentiation. Samples were collected at Days 0, 4, 7, 14, 21, and 28 for proteomic (n = 3 per time point) and phosphoproteomic analysis (n = 3 per time point). (B-C) Schematic overview of the workflows for automated total proteomics (B) and phosphoproteomics (C), including key steps of sample preparation, peptide enrichment, and data analysis pipelines. (D) Representative live-cell imaging of neurite outgrowth at different time points, showing stained soma (red) and dendrites (blue). The scale bar is 200µm. Total neurite length (mm/mm²) was quantified every day and was presented as mean ± SD. (E) Identifications of peptide, protein, phosphopeptide and phosphoprotein across different time points. Each point represents the mean count across replicates. Statistical significance was assessed using two-way ANOVA with Tukey’s post hoc test (*P < 0.05, **P < 0.01, ***P < 0.001), error bars representing mean ± SD. (F) Quantification of pluripotency markers (left) and mature neuronal markers (right). Log_2_ fold change (DN/D0) was calculated relative to Day 0 (iPSC) with error bars representing mean ± SD. (G) Principal component analysis (PCA) of proteome (left) and phosphopeptide (right) data colored by different days.

To optimize phosphopeptide enrichment efficiency, we systematically evaluated various types of magnetic beads and peptide input amounts. Mixed Ti-IMAC HP and Zr-IMAC HP beads (MagReSyn, 1:1 ratio) with a 200 μg protein input achieved the highest phosphopeptide identification (>15,000) and the best reproducibility (Figure S1A-C). Our optimized workflows demonstrated high coverage and reproducibility in quantifying proteins and phosphopeptides (Figure 1E, Table S1-2). Overall, we quantified 10,137 protein groups and 23,168 phosphorylated peptides with low missing values (Figure 1E, S1D-E) in both datasets. Identified peptides and proteins were relatively stable across all time points. In contrast, identified phosphopeptides increased significantly over time (p-value < 0.05), suggesting an elevation in phosphorylation-dependent signaling as neurons mature (Figure 1E). In our dataset, 44.7% of the total proteins identified have at least one phosphosite (Figure S1F), primarily serine (80.12%) followed by threonine (15.7%) and tyrosine residues (4.18%) (Figure S1G). Based on the longitudinal proteomics data, we evaluated the known biomarkers of neuronal maturation in KOLF2.1J-derived neurons. As expected, the protein expression of pluripotency markers POU5F1 (also known as OCT4) and SOX2 were significantly decreased within the first 14 days (Figure 1F). In contrast, mature neuronal markers, including NEFM, and MAP2 were rapidly upregulated within the first 7 days followed by a plateau during the later stages of differentiation (D14-D28). Additionally, RBFOX3, and NEFH and MAPT (also known as Tau) showed a steady increase in expression throughout differentiation (Figure 1F). Multiple markers were further validated the expression and localization via immunostaining (Figure S2A). Additionally, from the proteomics data, we recovered multiple key markers associated with synapses, cortical neurons, glutamatergic neurons, cholinergic neurons, and GABAergic neurons (Figure S2B). Principal component analysis (PCA) of the proteomic and phosphoproteomic data revealed high reproducibility in biological replicates and clear temporal separation across differentiation time points (Figure 1G). Compared to the proteome, the phosphoproteomic profile that had a higher number of features exhibited more distinct molecular signatures associated with neuronal maturation at later stages (D14-D28).

### Temporal clustering reveals dynamic regulation of biological pathways during neuronal differentiation

To investigate the temporal regulation of proteomic and phosphoproteomic dynamics during neuronal differentiation, we applied Mfuzz clustering analysis in both datasets, grouping proteins and phosphopeptides into eight clusters based on their temporal expression trajectories (Figure 2). Gene Ontology (GO) analysis was then performed to identify the top enriched pathways within each cluster. Comparing the clustering patterns of proteomics (Figure 2A) and phosphoproteomics (Figure 2B) revealed both shared and distinct regulatory mechanisms. In Clusters 1 and 2, proteins exhibited continuous upregulation throughout differentiation, focusing on pathways involved in axon and dendrite development, neuron projection organization, and synaptic signaling. However, the phosphorylation of proteins associated with similar functions plateaued early, as observed in Clusters 1–3 of Figure 2B. This suggests that phosphorylation may play a key role in initial regulatory mechanisms during neuronal differentiation. Notably, we observed that mitochondrial ATP synthesis and oxidative phosphorylation pathways showed progressive upregulation in the proteome (Cluster 1), supporting studies showing that mitochondrial metabolic activity may temporally accelerate neuronal development^24^.

**Figure 2.**
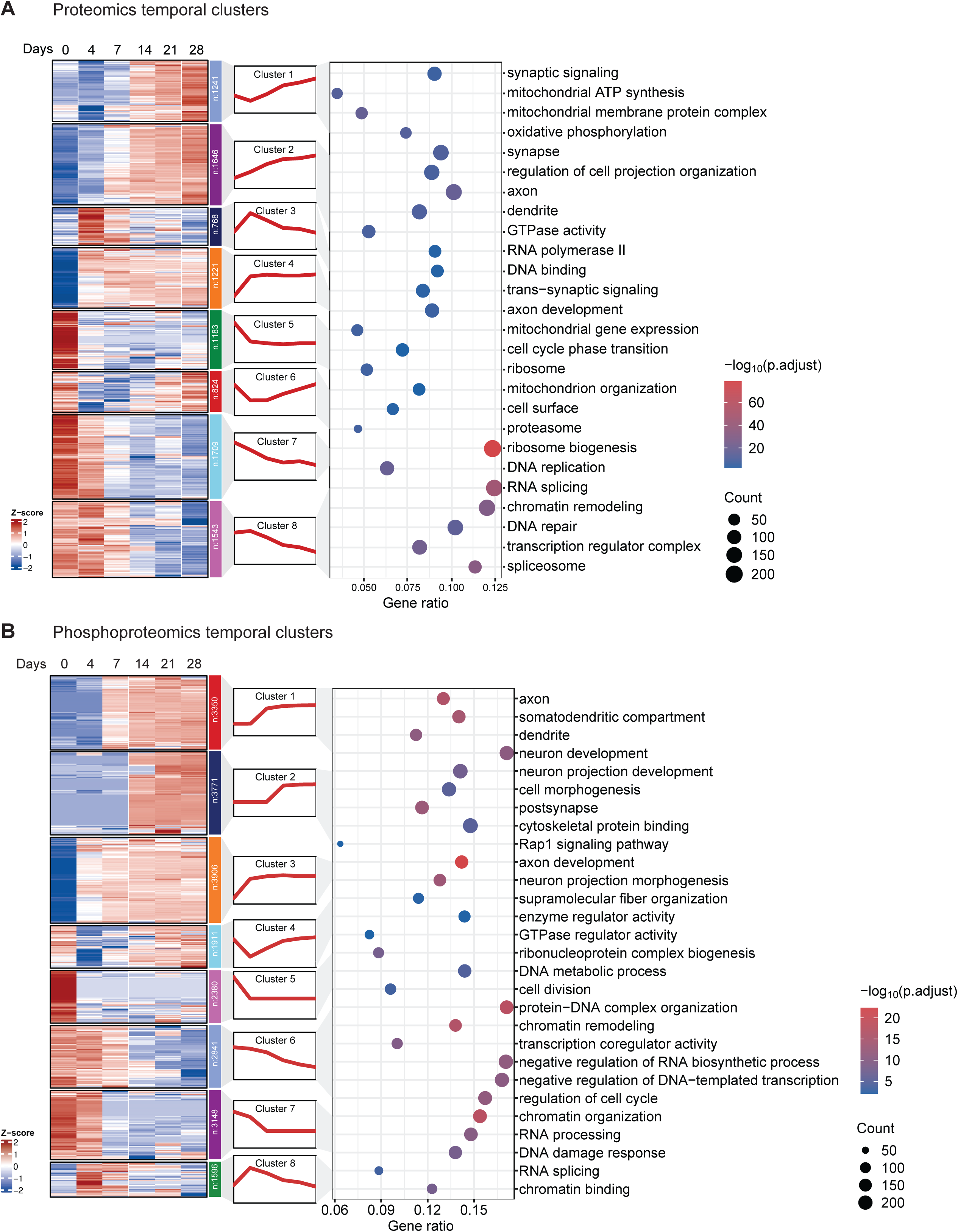
Clustering and pathway enrichment of the trajectories of proteome and phosphoproteome differentiation. (A) Proteome temporal clustering and GO enrichment analysis. The heatmap illustrated the eight temporal clusters using Mfuzz analysis, with each cluster representing proteins exhibiting similar expression trends. Z-scores indicated scaled relative expression levels. The bubble plot showed the GO enrichment analysis of each cluster, highlighting the top-enriched biological processes. (B) Phosphoproteome temporal clustering and GO enrichment analysis. The same Mfuzz approach was applied to the phosphoproteome data, grouping phosphoproteins into eight clusters based on their phosphopeptide quantitative dynamics across differentiation stages. GO enrichment analysis was performed based on the corresponding proteins of the phosphopeptides. X-axis represents the gene ratio, bubble size represents the gene count, and color intensity indicates the - log10 adjusted p-value, FDR<0.05.

We also found the transient upregulation of RNA polymerase II-associated proteins and DNA-binding factors in Cluster 3, peaking at D4, indicating a critical transcriptomic switch in early differentiation. Similarly, Cluster 6 in the phosphoproteome was enriched for chromatin remodeling and transcriptional regulation, suggesting that phosphorylation-driven reprogramming of gene expression facilitates the transition from pluripotency to a neuronal state. Additionally, GTPase-related phosphorylation events showed a biphasic regulation, with phosphorylation suppressed at D4 followed by a gradual activation until D28. This trend aligns with previous studies demonstrating the role of GTPases as key regulators of cytoskeletal remodeling and neurite outgrowth^25^. The co-enrichment of supramolecular fiber organization pathways with GTPase phosphorylation events in cluster 4 highlighted the phosphorylation-dependent mechanism for controlling cytoskeletal dynamics during axon extension and neuronal maturation.

Finally, pathways exhibiting progressive downregulation during differentiation included DNA replication and RNA splicing at the proteome level, as well as cell cycle regulation, chromatin organization, and RNA processing at the phosphoproteome level. These changes reflect the transition from a proliferative to a post-mitotic neuronal state. This analysis provided a deep exploration of the temporal proteomic changes during neuronal differentiation, with phosphoproteomics data offering additional granularity by characterizing the phosphorylation-dependent regulation of these processes.

### Functional pathways reveal the essential phosphosites for neuronal differentiation

To further investigate the roles of key proteins and phosphosites during neuronal maturation, we combined both omics datasets to provide a multilayer resource for functional validations. We further analyzed the protein list from the Gene Ontology (GO) database and overlapped it with our identified proteins and phosphosites. We generated heatmaps to depict temporal trends, with both proteome and phosphoproteome data z-scaled for comparative analysis. We focused on three key pathways: mitochondrial membrane organization (GO:0007006), oxidative phosphorylation (GO:0006119), and the tricarboxylic acid (TCA) cycle (GO:0006099). Our analysis identified six phosphorylated proteins involved in oxidative phosphorylation^26^ and six in the TCA cycle^27^. The TCA pathway exhibited synchronous expression patterns across both datasets, demonstrating concerted upregulation throughout differentiation. Interestingly, while the oxidative phosphorylation pathway showed overall upregulation at the protein level, some of its phosphosites displayed discordant trends. For instance, ATP5MG T21 and NDUFV3 S105 peaked during the pluripotency stage but were attenuated in mature neurons, in contrast to expression of the respective underlying proteins. Mitochondrial membrane proteins, such as those in the CHCHD and ATF2 families, exhibited minimal changes at the phosphoproteome level compared to their protein levels. However, kinases associated with mitochondrial function, such as GSK3A and GSK3B, showed robust phosphorylation up-regulation during later stages of neuronal differentiation (Figure 3A).

**Figure 3.**
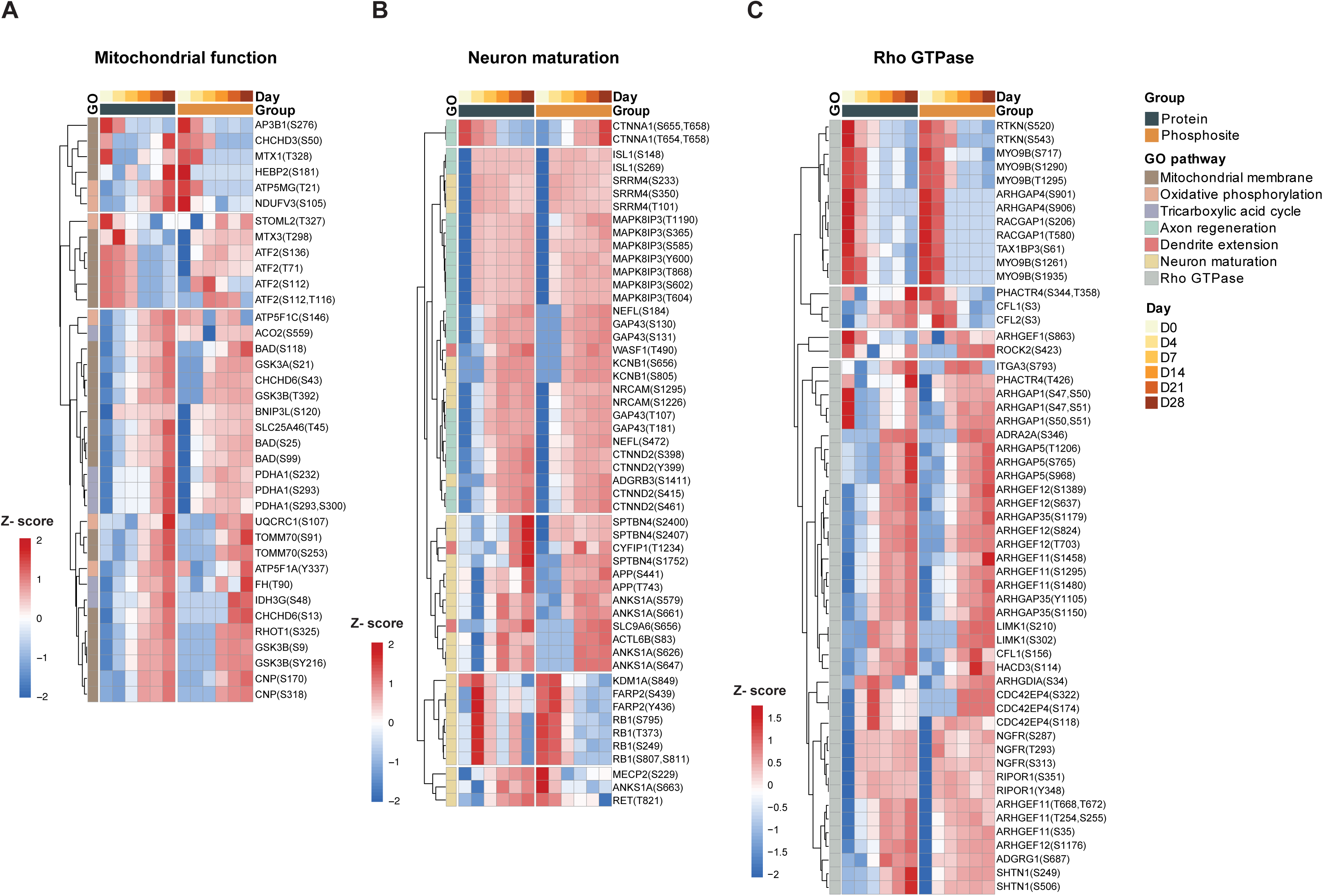
Dynamic trends in total protein and phosphosite levels within functional pathways critical for neuronal differentiation. (A) Heatmap showing the quantified key protein and phosphosites related to mitochondrial function pathways. (B) Heatmap showing the quantified key protein and phosphosites related to neuronal maturation pathways. (C) Heatmap showing the quantified key protein and phosphosites related to Rho GTPase pathways. Protein and phosphosite expression levels were z-scaled separately. Complete linkage cluster were applied based on phosphosites data.

To capture the pathways essential for neuronal maturation, we focused on axon regeneration (GO:0031103), dendrite extension (GO:0097484), and neuron maturation (GO:0042551). As expected, the majority of phosphosites and their corresponding proteins were upregulated early in differentiation, reaching a plateau shortly after induction. Notably, CTNNA1, a protein responsible for progenitor cell proliferation^28^, exhibited gradual downregulation at the protein level while its phosphosites increased. The axon regeneration pathway showed consistent trends between total proteomics and phosphoproteomics, with multiple phosphosites identified in critical axon-related proteins such as ISL1, MAPK8IP3, NEFL, GAP43, and CTNND2^29–32^. Similarly, the neuron maturation pathway revealed early phosphorylation events, particularly in proteins such as SPTBN4 which maintains axonal cytoskeleton integrity^33^; APP which plays a key role in synapse formation^34^; and RB1which is a regulator of cell cycle arrest^35^, that exhibited highly regulated phosphorylation at the iPSC and D4 stages (Figure 3B). Additionally, small GTPase proteins from the ARHGAP and ARHGEF families displayed more dynamic phosphorylation changes than protein level changes, with inhibition observed at D4, potentially driving neuronal maturation (Figure 3C).

Microtubule dynamic changes, while slightly delayed compared to protein dynamics, showed pronounced changes in later stages, particularly in MAPT and MAP1B (Figure S3A). Spearman correlation analysis revealed that although more than half of the proteins exhibited positive correlations with their phosphosites, a significant subset showed negative correlations (Figure S4B). Further analysis of the pathways with negative correlations highlighted RNA transport and localization as a key pathway of interest, Notably, proteins such as AGFG1, TPR, SRSF, MUP, and HNRNPA (Figure S4C) were implicated, suggesting that the function of RNA-binding and transport proteins can be regulated through phosphorylation-dependent processes^36–38^.

### Kinome maps reveal the kinase dynamics for neuronal differentiation

To investigate kinome dynamics, we performed kinase-substrate enrichment analysis (KSEA) on significantly altered phosphosites to identify early stage (D7 vs. D0) changes (Figure 4A) and late stage (D28 vs. D7) changes (Figure 4B) of differentiation. KSEA infers kinase activity by assessing phosphorylation changes in their known substrates. Our data suggested that each kinase family exhibited distinct variations during the differentiation. Importantly, the tyrosine kinase-like (TKL) family was preferentially activated with key kinases such as RAF, ARAF, JAK2, and RET showing early upregulation within the first 7 days. This observation is consistent with previous studies demonstrating that RAF kinases regulate GTPase signaling and JAK/STAT pathways^39–41^, both of which are critical for presynaptic formation. PRKAA2, a component of the AMP-activated protein kinase (AMPK), displayed the most significant kinase enrichment across multiple time points (Figure S4). Additionally, Rho-associated coiled-coil containing protein kinase 1 (ROCK1) and ROCK2 expression, key regulators of Rho GTPase activity, aligned with their phosphosites trend observed in our dataset (Figure 3C) and are known to drive neuronal maturation and synaptic plasticity^25^. At later stages of differentiation, kinase activity shifted toward the tyrosine kinase (TK) family, with a significant increase in FYN, ALK, and HCK. Conversely, CDK family kinases, particularly CDK1, CDK2, and CDK7, were progressively downregulated throughout differentiation (Figure 4C, S4).

**Figure 4.**
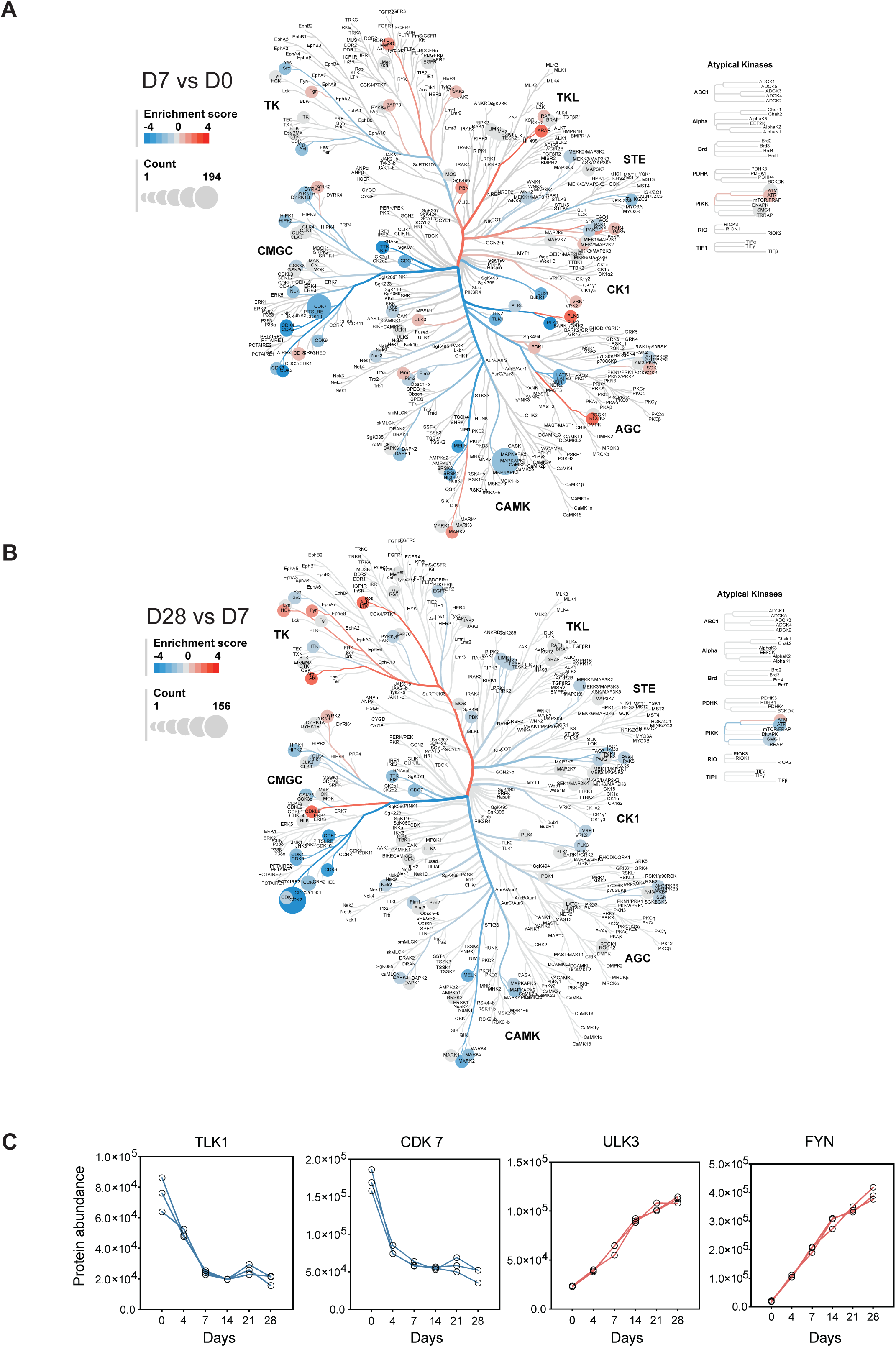
Kinome map reveals the dynamic changes during the neuronal differentiation. (A) Kinase substrate enrichment analysis (KSEA) between early phase Day 7 iNeuron vs iPSC (Day 0). (B) Kinase substrate enrichment analysis (KSEA) between late phase Day 28 vs Day 7. Branch and node color represent the enrichment score. Node size represents the counts of substrates. (C) Temporal expression profiles of representative kinase protein abundance.

In addition to performing KSEA, which provides insights into kinase activity through kinase-substrate interactions, we also analyzed the protein abundance of kinases in the proteomics dataset to validate the KSEA results. For example, TLK1 and CDK7, critical kinases that regulate the cell cycle and chromatin remodeling, exhibited strong and consistent downregulation. In contrast, ULK3 and FYN, key regulators of neuronal development and potential therapeutic targets for neurodevelopmental disorders, were upregulated at the protein level. Together, these findings highlighted a dynamic and stage-specific shift of kinase activity during neuronal differentiation.

## Discussion

Previous studies of neuronal differentiation have been mainly limited to transcriptomic and proteomic analyses. Here, we sought to add additional insights at the PTMs level underlying the transition from iPSC pluripotency exit to neuron maturation. Utilizing the KOLF2.1J iPSC line and a robust differentiation protocol, we established an automated protein and phosphopeptide enrichment pipeline to achieve in-depth proteome and phosphoproteome profiling at matched time points throughout neuronal differentiation. Our study highlighted the activation and inhibition of key pathways across distinct developmental stages, including the pluripotency phase, immature stage, and mature neuronal stage, providing a systematic roadmap of molecular changes during neuronal differentiation (Figure 5). To easily explore our results, we made all proteomic and phosphoproteomics data available for further data visualization in a user-friendly web app. Researchers will find this as a useful resource for visualizing protein abundance and phosphosite levels at different neuronal maturation stages.

**Figure 5.**
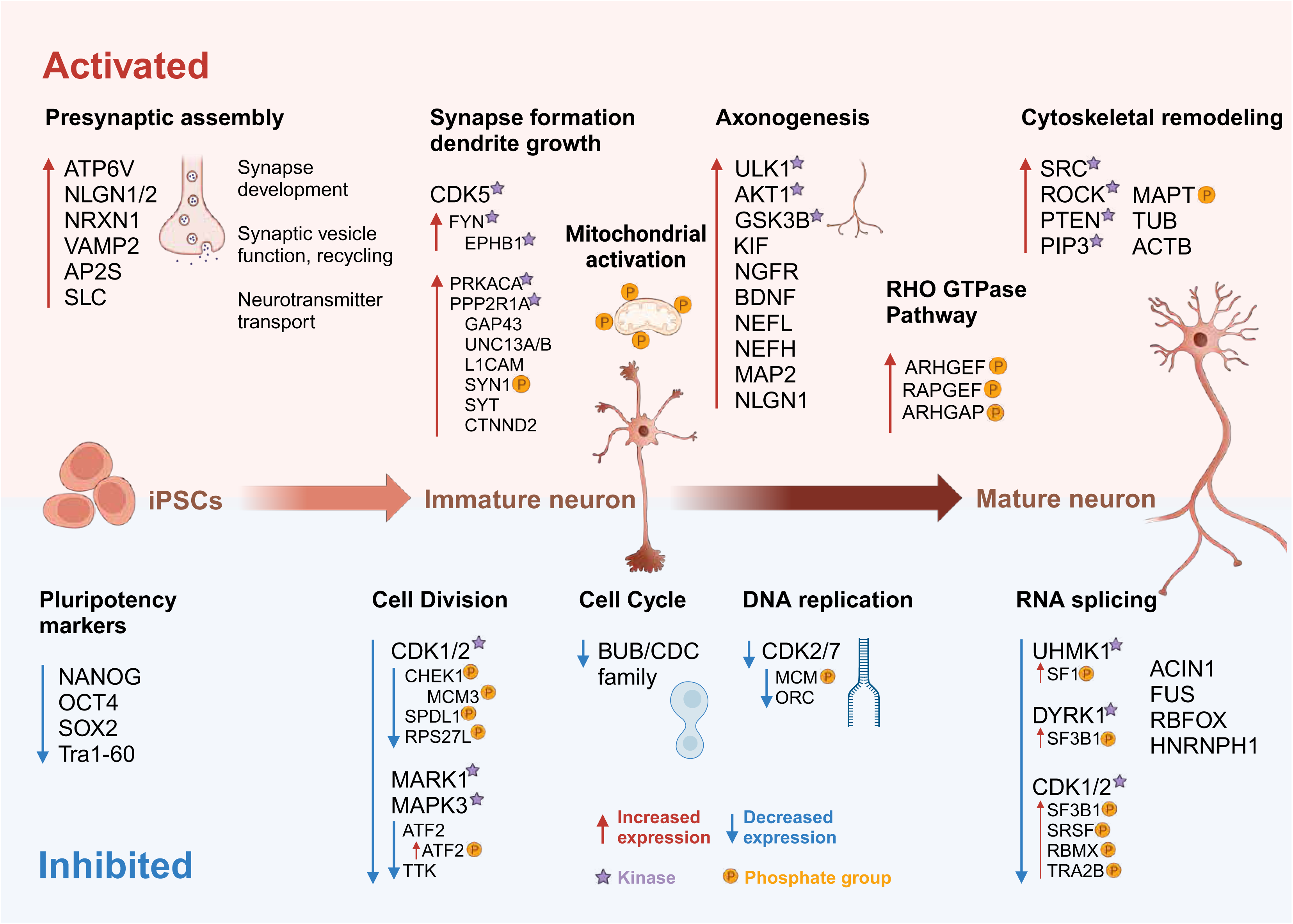
Roadmap of neuronal differentiation. Summary of key proteins, phosphorylated protein, kinases, and signaling pathways distinguish regulated during the KOLF2.1J neuron differentiation.

Proteins associated with cell proliferation exhibited high expression during early differentiation. However, the decline of traditional stem cell markers such as SOX2 and OCT4, occurred at varying rates (Figure 1F), making them insufficient for defining pluripotency exit. Phosphoproteomic data revealed a marked decline in ribosome biogenesis and chromatin remodeling. This suggested that phosphorylation events could serve as additional indicators for assessing stem cell maintenance and optimizing differentiation protocols. Furthermore, kinase activity analysis showed that the CDK family, key regulators of the cell cycle, was highly active in iPSCs but exhibited sustained downregulation during differentiation. We provided evidence that CDK-substrate interactions contribute to the regulation of pluripotency exit via our comprehensive evaluation of early differentiation fidelity at the protein and phosphorylation level. Mitochondrial dynamics and metabolism exhibited temporal changes throughout neuronal development. The expression of mitochondrial proteins was relatively low during the iPSC and early differentiation stages but gradually increased as neurons matured. This trend aligns with previous studies^24,42^, which suggests that enhanced mitochondrial metabolism in postmitotic neurons accelerates maturation. We observed that the majority of proteins associated with the TCA cycle and oxidative phosphorylation followed by similar trends in iPSC-derived neurons. Furthermore, we identified key mitochondrial phosphorylation events that are tightly correlated with neuronal maturation.

Neurons are highly polarized cells, and our data revealed robust activation of axon and dendrite pathways during the intermediate stage of differentiation. Microtubules, which serve as critical tracks for intracellular transport to axons and dendrites^20,43^, exhibited extensive phosphorylation events throughout neuronal development. Notably, we observed significant phosphorylation of MAP1B, particularly at the late stage, suggesting its role in maintaining neuronal polarity and supporting neuronal maturation. Interestingly, we also identified highly active phosphorylation of MAPT (also known as Tau), a well-known marker of Alzheimer’s disease pathology when hyperphosphorylated^44^. The phosphorylation of Tau at specific sites during neuronal maturation may reflect its physiological role in microtubule stabilization, while aberrant phosphorylation at these sites in disease states could contribute to tauopathies. Additionally, Rho GTPases, which act as molecular switches coordinating actin and microtubule dynamics, exhibited numerous phosphosites in our dataset. Many of these sites have not been previously reported, providing a valuable resource for research community.

To our knowledge, this is the first study to comprehensively characterize kinome dynamics during neuronal differentiation. Our KSEA study enables stage-specific identification of kinome-wide patterns, offering a foundation for future studies exploring kinase-substrate relationships in neuronal differentiation and disease contexts.

In conclusion, our study provided a comprehensive temporal map of the proteome, phosphoproteome, and kinome during the development of KOLF2.1J-derived neurons, shedding light on the molecular mechanisms underlying neuronal differentiation and maturation. These findings not only enhance our understanding of neuronal development but also offer a valuable resource for investigating the molecular basis of neurodevelopmental and neurodegenerative disorders using iPSC-derived neuronal models.

### Limitations of the study

This study aimed to achieve unbiased phosphopeptide enrichment. However, quantifying phosphosites with low stoichiometry remains challenging. Future advancements in single-cell proteomics and phosphoproteomics could provide deeper insights into the heterogeneity of neuronal populations and their developmental trajectories. The phosphosites and kinases described in this study merit further functional validation to elucidate their molecular mechanisms. Additionally, this study did not assess the variability of the KOLF2.1J line in comparison to other iPSC lines. There is a lack of integration of phosphoproteomic data with transcriptomics or other PTM datasets; such integration could offer a more comprehensive view of regulatory networks governing neuronal differentiation.

## STAR Methods

### Key resources table

| REAGENT or RESOURCE | SOURCE | IDENTIFIER |
| --- | --- | --- |
| **Chemicals, peptides, and recombinant proteins** |  |  |
| PB-TO-oNGN2 | Addgene | Cat# 198397 |
| EFa1-Transposase | Addgene | Cat# 172116 |
| Puromycin | Sigma-Aldrich | Cat# P9620 |
| Accutase | Thermo Fisher | Cat# A111050 |
| Matrigel | Corning | Cat# 354277 |
| DMEM/F12-HEPES | Thermo Fisher | Cat# 11330032 |
| N2 Supplement | Thermo Fisher | Cat# 17502048 |
| Non-Essential Amino Acids | Thermo Fisher | Cat# 11140-050 |
| Glutamax | Thermo Fisher | Cat# 35050-061 |
| Chroman 1 | MedChemExpress | Cat# HY-15392 |
| Ammonium bicarbonate (>99%) | Millipore Sigma | Cat# A6141 |
| Benzonase Nuclease, ultrapure | Millipore Sigma | Cat# E8263 |
| Complete protease inhibitor | Millipore Sigma | Cat# 11697498001 |
| Ethyl Alcohol, Anhydrous, 200 Proof | Electron Microscopy Sciences | Cat# 15055 |
| Glycerol, ultra pure | MP Biomedicals | Cat# 800687 |
| NP-40 | Thermo Scientific | Cat# 85124 |
| Acetonitrile (ACN), LC-MS Grade | Thermo Scientific | Cat# TS-51101 |
| Dithiothreitol | Thermo Scientific | Cat# 20290 |
| Glycolic acid | Thermo Scientific | Cat# 154511000 |
| Iodoacetamide, Single-Use | Thermo Scientific | Cat# A39271 |
| Water, LC-MS Grade | Thermo Scientific | Cat# 51140 |
| Trifluoroacetic Acid (TFA), LC-MS Grade | Thermo Scientific | Cat# 85183 |
| Formic Acid, LC-MS Grade | Thermo Scientific | Cat# 28905 |
| Sodium deoxycholate (>97%) | Millipore Sigma | Cat# D6750 |
| Magnetic carboxylate modified particles | Millipore Sigma | Cat# GE65152105050250 |
| PureCube Fe-NTA MagBeads | Cube biotech | Cat# 31501-Ti |
| High Select™ Phosphopeptide Enrichment Kits | Thermo Scientific | Cat# A32993 |
| Ti-IMAC HP beads | Resyn Biosciences | MR-THP002 |
| Zr-IMAC HP beads | Resyn Biosciences | MR-ZHP002 |
| Ammonium hydroxide solution | Millipore Sigma | Cat# 1336-21-6 |
| Triton X-100 | Millipore Sigma | Cat# X100 |
| Trypsin / Lys-C Mix, Mass Spec Grade | Promega | Cat# V5073 |
| TWEEN 20 | Millipore Sigma | Cat# P9416 |
| 0.5M EDTA, pH 8.0 | Thermo Scientific | Cat# 15575020 |
| 1M Tris-HCl, pH 8.0 | Thermo Scientific | Cat# 15568025 |
| 5M NaCl | Thermo Scientific | Cat# 24740011 |
| SDS Solution, 10% | Thermo Scientific | Cat# 15553027 |
| Bovine Serum Albumin | Thermo Scientific | Cat# A9418 |
| Saponin | Sigma-Aldrich | Cat# 84510 |
| Essential 8 medium | Thermo Scientific | Cat# A1517001 |
| Dulbecco's Phosphate-Buffered Saline (DPBS) | Life Technologies Corporation | Cat# 14190250 |
| Cell Culture Grade Water | Quality Biological | Cat#118-162-151 |
| Chroman I | MedChem Express | Cat# HY-15392 |
| Knockout DMEM/F12 | Thermo Scientific | Cat# 12660012 |
| N-2 Supplement | Thermo Scientific | Cat# 17502001 |
| Paraformaldehyde | EMS | Cat# 15710 |
| ROCK inhibitor Y-27632 | SelleckChem | Cat# S1049 |
| Doxycycline | Sigma-Aldrich | Cat# D9891 |
| poly-L-ornithine hydrobromide | Sigma-Aldrich | Cat# P3655 |
| Brainphys neuronal maturation medium | Stemcell Technologies | Cat# 5790 |
| N21-MAX Supplement | R&D Systems | Cat# AR008 |
| GDNF | PeproTech | Cat# 450-10 |
| BDNF | PeproTech | Cat# 450-02 |
| NT3 | PeproTech | Cat# 450-03 |
| Laminin | R&D Systems | Cat# 3446-005-01 |
| 4',6-diamidino-2-phenylindole (DAPI) | Thermo Fisher | Cat# 62248 |
| PBS | Lonza | Cat# 17-516F/12 |
| Dimethyl sulfoxide (DMSO) LC-MS Grade | Thermo Scientific | Cat# 85190 |
| 5-Fluoro-2'-deoxyuridine | Sigma-Aldrich | Cat# F0503 |
| Uridine | Sigma-Aldrich | Cat# U3003 |
| Sep-Pak C18 96-well Plate | Waters | Cat# 186003966 |
| **Critical commercial assays** |  |  |
| Lipofectamine Stem Transfection Reagent | Thermo Fisher | Cat# STEM00008 |
| Detergent Compatible Protein Assay | Bio-Rad | Cat# 5000111 |
| **Deposited data** |  |  |
| MS-based proteomic raw data files | ProteomeXchange Consortium | PXD061320 |
| **Cell line** |  |  |
| KOLF2.1J stem cell line | The Jackson Laboratory | JIPSC1000 |
| **Software and algorithms** |  |  |
| Spectronaut | Biognosys | Version 19 |
| R | The R Project | Version 4.3 |
| ProtPipe | Ziyi et.al., 2024 |  |
| GraphPad Prism | GraphPad Software, San Diego, California USA | Version 10 |
| Adobe Illustrator | Adobe | Version 29.3 |
| **Other** |  |  |
| EASY-Spray nano HPLC column | Thermo Scientific | Cat# ES903 |
| Acclaim PepMap trap column | Thermo Scientific | Cat# 164535 |
| Matrix 0.5mL ScrewTop Tubes | Thermo Scientific | Cat# 3744 |
| 96 well polypropylene deep well plate | Corning | Cat# CLS3960 |

### Resource availability

#### Lead contact

Further information and requests for resources and reagents should be directed to and will be fulfilled by the lead contact, Yue Andy Qi.

### Materials availability

This study did not generate new unique reagents.

### Experimental model and subject details

#### Cell lines

In this study, we used the KOLF2.1J line derived from a caucasian male, obtained from a reference parental line of the iNDI project (The Jackson Laboratory, https://www.jax.org/jax-mice-and-services/ipsc). Material transfer agreement between the Jackson Laboratory and the Center for Alzheimer’s and Related Dementias at the NIH was obtained prior to conducting any experiment.

### Data and code availability

The mass spectrometry proteomics data have been deposited to the ProteomeXchange Consortium via the PRIDE^45^ partner repository. The accession number is listed in the key resources table.

This paper does not report original code.

Any additional information required to reanalyze the data reported in this paper is available from the lead contact upon request

### Method details

#### Neuronal differentiation of human iPSC

We utilized the iNDI reference parental iPSC KOLF2.1J line, and neurons were differentiated as previously described^7,23^ with some modifications (https://dx.doi.org/10.17504/protocols.io.8epv5zyn5v1b/v1). Briefly, the KOLF2.1J cells were transfected with a PB-TO-oNGN2 vector and EFa1-Transposase with a 1:3 ratio (transposase: vector) and a total of 3µg of DNA mix, using Lipofectamine Stem Transfection Reagent followed by selection after 24-48 hrs. with 1 to 8 µg/ml puromycin for a minimum of 72 hours up to two weeks. The selected iPSCs with a stably integrated human NGN2 were singularized using Accutase and plated at 2.5x10^5^ per well into a Matrigel-coated (1:100) 6 well plates for 4 days with Neuronal Induction Media (NIM) containing DMEM/F12-HEPES, 1X N2 Supplement, 1X Non-Essential Amino Acids, 1X Glutamax, 50 nM Chroman 1 and 2 µg/mL Doxycycline. On D4, immature neurons were dissociated using Accutase and replated at 1.5 x10^6^ cells per well into a poly-L-ornithine coated 6 well plate using Neuronal Maturation Media (NMM) for D4 to D7, containing DMEM/F12-HEPES, Brainphys Neuronal Medium (1:1), 1X N21-MAX Supplement, 10 ng/mL GDNF, 10 ng/mL BDNF, 10 ng/mL NT3, 1 µg/mL Laminin, 2 µg/mL Doxycycline, 5-Fluoro-2′-deoxyuridine 1 µM and Uridine. For D10 to D28 medium was replaced using NMM containing Brainphys Neuronal Media only plus all the supplements used at day 4 to 7. For neuronal maintenance after day 4, half-medium changes were performed every 3-4 days until day 28.

#### Live-cell imaging for neurite outgrowth

KOLF2.1J cells with stably integrated human NGN2 were transduced with a lentivirus expressing cytosolic mScarlet^46^ (MK-EF1a-mScarlet) to identify neurites. The course of the neuronal differentiation was recorded by an Incucyte S3 Live-Cell Analysis System with a 20X objective every 24 h at 37°C for 28 days and phase and fluorescent images were acquired for every time point. The neurite outgrowth was analyzed with the NeuroTrack Incucyte software module.

#### Immunofluorescence and imaging

For the immunofluorescence imaging of neuron and axon markers, phase and fluorescent images were acquired for every time point. Cells were fixed in 4% Paraformaldehyde for 10 min at room temperature and washed with 1x DPBS three times for 5 min each. Primary and secondary antibody dilutions were prepared in 1x DPBS supplemented with 3% Bovine Serum Albumin and 0.2% Saponin. Fixed cells were incubated with primary antibodies overnight at 4°C, followed by 3 time washes with 1x DPBS. Cells were then incubated with secondary antibodies for 1 h at room temperature in the dark. Last, DAPI stain (1:3000 diluted in 1x DPBS) was applied for 30 min at room temperature and followed by 2 washes with 1x DPBS. Images were acquired on the Nikon Eclipse Ti2 Confocal Microscope, using a 60X Water Immersion lens. Images were analyzed using the NIS-Elements AR software.

#### Automated sample preparation for proteomics

Sample preparation for proteomics was conducted on a fully automated workflow based on our previous report^47^. Briefly, cells were washed thoroughly with ice-cold PBS and 200 μL lysis buffer were added directly to each well of a 6-well plate. The lysis buffer consisted of 50 mM Tris-HCI (pH = 8.0), 50 mM NaCl, 1% SDS, 1% triton X-100, 1% NP-40, 1% tween-20, 1% glycerol, 1% sodium deoxycholate (wt/vol), 5 mM EDTA (pH = 8.0), 5mM dithiothreitol (DTT), 5KU benzonase, and 1× complete protease inhibitor. The cell lysates were scraped off the plate surface and harvested into a 96 deep well plate tube. The lysates were incubated at 65°C for 30 minutes on a Thermomixer allowing for protein denature and followed by alkylation using 10 mM iodoacetamide in the dark. Protein concentrations were normalized to 0.2 mg/mL for total proteomics analysis using automated Detergent Compatible Protein Assay. The protein enrichment and on-bead tryptic digestion were conducted using a KingFisher APEX robotic system. Proteins were digested with Trypsin/Lys-C mix in 50 mM ammonium bicarbonate at 37°C for 16 hours. Following digestion, peptides were vacuum-dried and reconstituted in 2% acetonitrile with 0.1% formic acid. 1μg of peptides was used for LC-MS/MS proteomic analysis.

#### Automated workflow of phosphopeptide enrichment

We evaluated 4 types of commercially available magnetic beads: PureCube Ti-NTA, PureCube Fe-NTA, MagReSyn Ti-IMAC HP and MagReSyn Zr-IMAC HP beads. High Select phosphopeptide manual enrichment kit was also used as the control (Figure 1SA). We optimized the phosphopeptetide enrichment method based on previous reports^48–50^. Briefly, dried tryptic peptides from 200μg of total protein digest were reconstituted in 100 μL loading buffer containing 80% acetonitrile (ACN), 5% trifluoroacetic acid (TFA), 0.1M glycolic acid (GA). A 1:1 ratio mixture of MagReSyn Ti-IMAC HP and Zr-IMAC HP beads was used for enrichment, with a bead-to-peptide ratio of 4:1. The beads were washed three times with the loading buffer prior to use. The enrichment procedure was fully automated on a KingFisher APEX robotic system. Peptides were incubated with beads for 20 minutes at medium mixing speed. The beads were then moved to three wash plates sequentially for 2 min each, first with loading buffer solvent (500μL), then wash solvent 1 (80% ACN + 1% TFA, 500 μL) and lastly by wash solvent 2 (10% ACN + 0.2% TFA, 500 μL). Phosphopeptides were eluted by incubation with 2% ammonium hydroxide (100 μL) for 10 minutes. The eluate was immediately acidified with 20μL of 10% formic acid. Desalting of enriched phosphopeptides was performed using solid-phase extraction (SPE) on Sep-Pak C18 96-well plates following the manufacturer protocol. The cleaned phosphopeptides were lyophilized and stored at -80°C. 1μg of phosphopeptides was injected for phosphoproteomic analysis.

#### Liquid chromatography and mass spectrometry

Total peptides and phosphopeptides were analyzed using an UltiMate 3000 nano-HPLC system (Thermo Scientific) coupled with an Orbitrap Eclipse mass spectrometer (Thermo Scientific). The peptides separation was performed on a ES903A nano column (75 μm × 500 mm, 2 μm C18 particle size) using an 80-minute linear gradient of 2% - 40% phase B (5% DMSO in 0.1% formic acid, in ACN). The column temperature was maintained at 60 °C and the flow rate was 300 nL/min. Data was acquired in data-independent acquisition (DIA) mode. The MS1 scan was set at a resolution of 120,000, with a standard AGC target, and the maximum injection time was set to auto. The MS2 scans covered a precursor mass range of 400–1000 m/z, using an isolation window of 8 m/z with 1 m/z overlap, resulting in a total of 75 windows per scan cycle. Fragmentation was performed using high-energy collisional dissociation (HCD) with a normalized collision energy of 30%. MS2 spectra were acquired at a resolution of 30,000, with a scan range of 145–1450 m/z, and the loop control was set to 3 seconds.

#### Data analysis for proteomics and phosphoproteomics

For database searching, we performed DIA database searches using Spectronaut (version 19, Biognosys) with the directDIA search strategy. We used the UniProt human proteome reference with reviewed genes (20,384 entries) as the library FASTA file. For proteomic analysis, we applied the factory default settings, while for phosphoproteomic analysis, we utilized the phospho PTM workflow. Trypsin/P and Lys-C were selected as the digestion enzymes. The digestion specificity was defined as specific, permitting a maximum of two missed cleavages per peptide. Carbamidomethylation of cysteine residues was set as a fixed modification. For variable modifications, oxidation of methionine and N-terminal acetylation were considered in both proteomic and phosphoproteomic analyses. Additionally, phosphorylation of serine, threonine, and tyrosine residues was included as a variable modification for phosphoproteomic analysis. The false discovery rate (FDR) thresholds for peptide-spectrum matches (PSMs), peptide, and protein groups were set to 0.01. To enhance data completeness while maintaining high-confident PTM localization, we employed a dual PTM localization probability cutoff strategy, as previously described^51^. Initially, a PTM localization probability cutoff of 0 was applied to export the “PTM000” dataset. Subsequently, the cutoff was set to 0.75, and the same export scheme was used to generate the “PTM075” dataset. The “PTM000” report was then filtered to retain only those precursors (EG.PrecursorId) that were present in the “PTM075” report. Cross-run normalization was enabled to account for systematic variations across different runs and no imputation was performed on the dataset. Unless explicitly stated, all other parameters were maintained as the default Spectronaut settings.

Further downstream analyses were performed using our in-house built informatic pipeline: ProtPipe^52^. Proteins or phosphopeptides were considered differentially expressed if the absolute value of the log₂ fold change was greater than 1 and an adjusted p-value < 0.05, with multiple testing correction applied using the Benjamini-Hochberg method. Functional enrichment analysis was performed using a q-value < 0.05 as significance thresholds. Heatmap was created using the R package “pheatmap”. GraphPad Prism (version10) was used to generate the histogram plots.

#### Mfuzz clustering analysis

We used the Mfuzz package^53^ in R, which applies fuzzy c-means clustering, allowing proteins or phosphosites to belong to multiple clusters with varying degrees of membership, with the optimal number of clusters (k = 8) determined based on centroid distance minimization and cluster stability. For visualization, we utilized the ClusterGVis package which enables interactive exploration and graphical representation of cluster trends. To functionally characterize each cluster, Gene Ontology (GO) enrichment analysis was performed using the clusterProfiler R package^54^, identifying significantly enriched pathways based on gene ratio, adjusted p-value (< 0.05), and gene count, providing insights into the dynamic regulation of biological processes during neuronal differentiation.

#### Kinase-substrate enrichment analysis (KSEA) and kinome visualization

To investigate the dynamics of kinase activity during neuronal differentiation, we performed Kinase-Substrate Enrichment Analysis (KSEA) using a curated kinase-substrate interaction database. KSEA estimates kinase activity by integrating phosphosite changes with known kinase-substrate relationships, enabling the identification of significantly regulated kinases. KSEA was performed using the ‘KSEAapp’ R package^55^ with a NetworKIN cutoff of 5. For this analysis, phosphosites were mapped to their corresponding kinases, and activity scores were calculated based on the log₂ fold change of phosphosite abundance. Significance was determined using adjusted p-values (< 0.05). We then used the Coral tool^56^ to annotate these kinases on a kinome tree. Nodes and branches in the kinome trees were color-coded based on enrichment scores from the KSEA results, allowing for the identification of key kinases involved in neuronal differentiation and revealing temporal patterns of kinase regulation.

#### Interactive web app construction

To visualize the temporal trends of proteome and phosphoproteome expression of KOLF2.1J-derived neurons, we generated a Shinyapp using the mean abundance and standard deviation across biological replicates for each protein and phosphosite. An interactive dropdown menu in the sidebar allows users to select one or multiple proteins of interest. Upon selection, the application dynamically updates to display available phosphosites associated with the chosen proteins. Users can select specific phosphosites to visualize their temporal expression patterns. The web app was developed using Shiny in RStudio (version 4.3.2) and deployed via shinyapps.io. available at https://niacard.shinyapps.io/Phosphoproteome/.

## Acknowledgement

This research was supported in part by the Intramural Research Program of the NIH, National Institute on Aging (NIA), National Institutes of Health, Department of Health and Human Services; Project number ZIAAG000535.

## Author contributions

Conceptualization, Y.H., Y.A.Q., Z.L., E.L.

Methodology, Y.H., E.L., I.K., J.C., M.S.

Data curation, Z.L., C.A.W.

Formal analysis, Z.L., C.A.W., J.E., N.C.

Investigation, Y.H., Z.L., E.L., D.R., M.S., B.J., J.E., N.C., I.K., C.W., S.K., M.A.N., P.N.3, Y.A.Q.

Resources, Z.L., C.A.W., L.F., A.B.S., M.E.W., M.R.C., Y.A.Q.

Writing – original draft, Y.H., Z.L.; Y.A.Q., E.L., M.S.

Writing – review & editing, all authors.

Visualization, Y.H., Z.L., J.E., N.C., P.J.

Supervision, L.F., A.B.S., M.E.W., P.N., M.R.C., Y.A.Q.

## Supplementary materials

**Figure S1:**
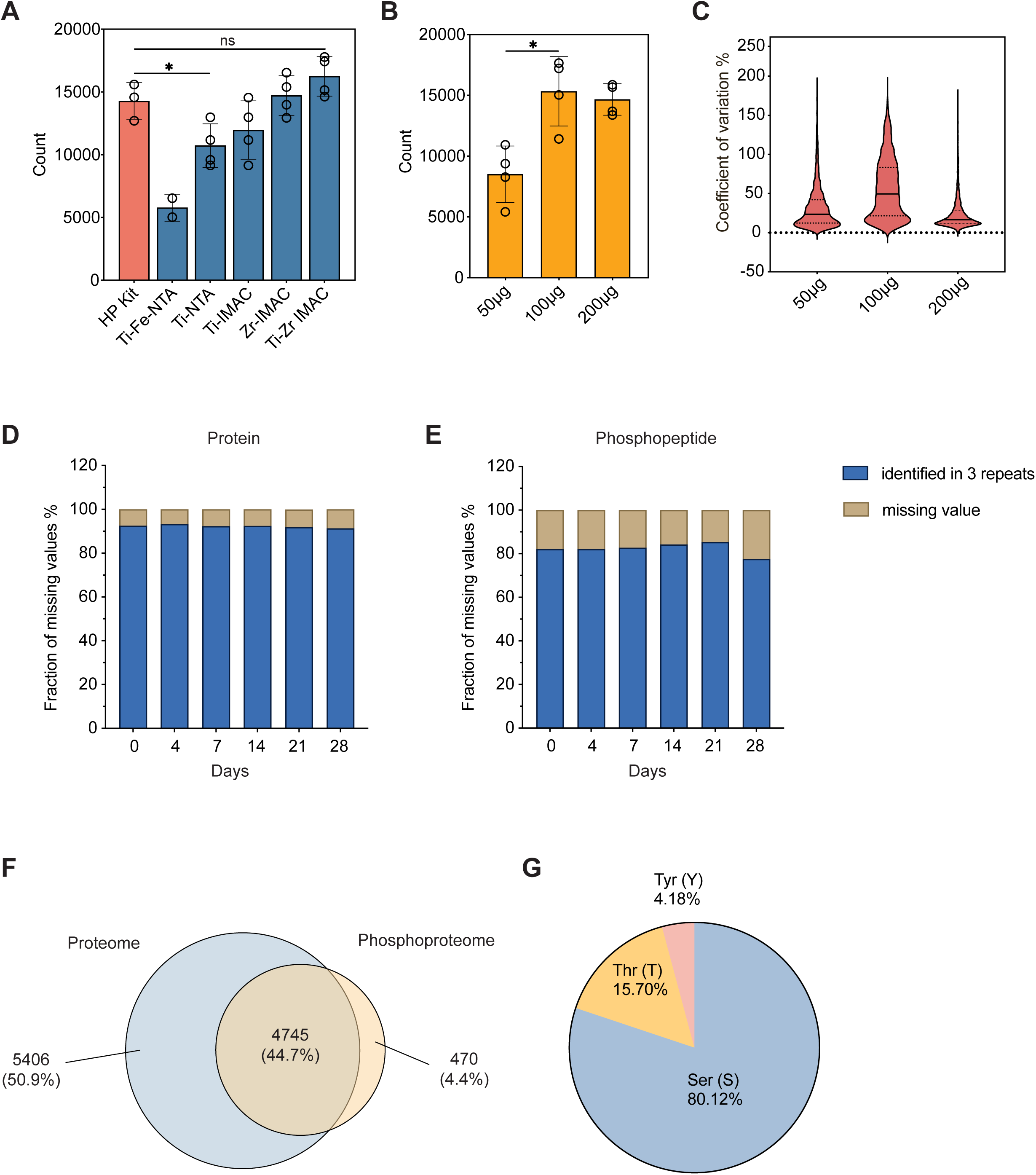
Optimization of phosphopeptide enrichment analysis. (A) Comparison of phosphopeptide enrichment efficiency across different commercial beads. Welch’s t test with *P < 0.05. (B) Effect of input protein amount on phosphopeptide identification. Welch’s t test with *P < 0.05. (C) Coefficient of variation (CV) analysis for phosphopeptide quantification across different input protein amounts. (D) Missing values of protein identification across the differentiation. (E) Missing values of phosphopeptide identification across the differentiation. (F) Venn diagram illustrating the overlap of the proteome and phosphoproteome overall identified in KOLF2.1J-derived neurons. (G) Distribution of phosphorylated residues.

**Figure S2:**
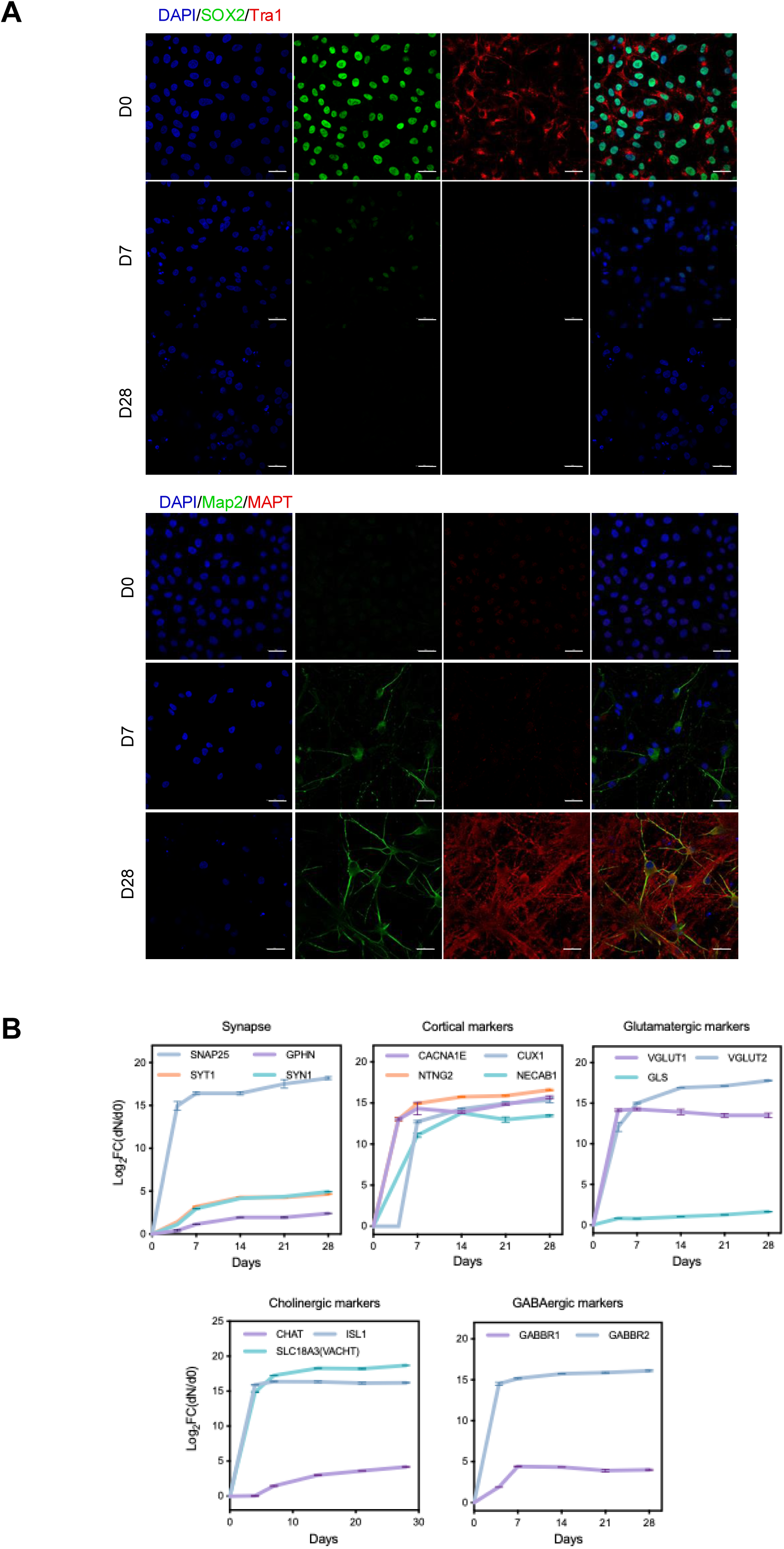
Characterization of neuronal markers. (A) Immunofluorescence staining of pluripotency and neuronal markers at different stages of differentiation. Scale bars represent 25 μm. (B) Protein quantification of different types of neuronal markers during differentiation. Log_2_ fold change (DN/D0) was calculated relative to Day 0 (iPSC) with error bars representing mean ± SD.

**Figure S3:**
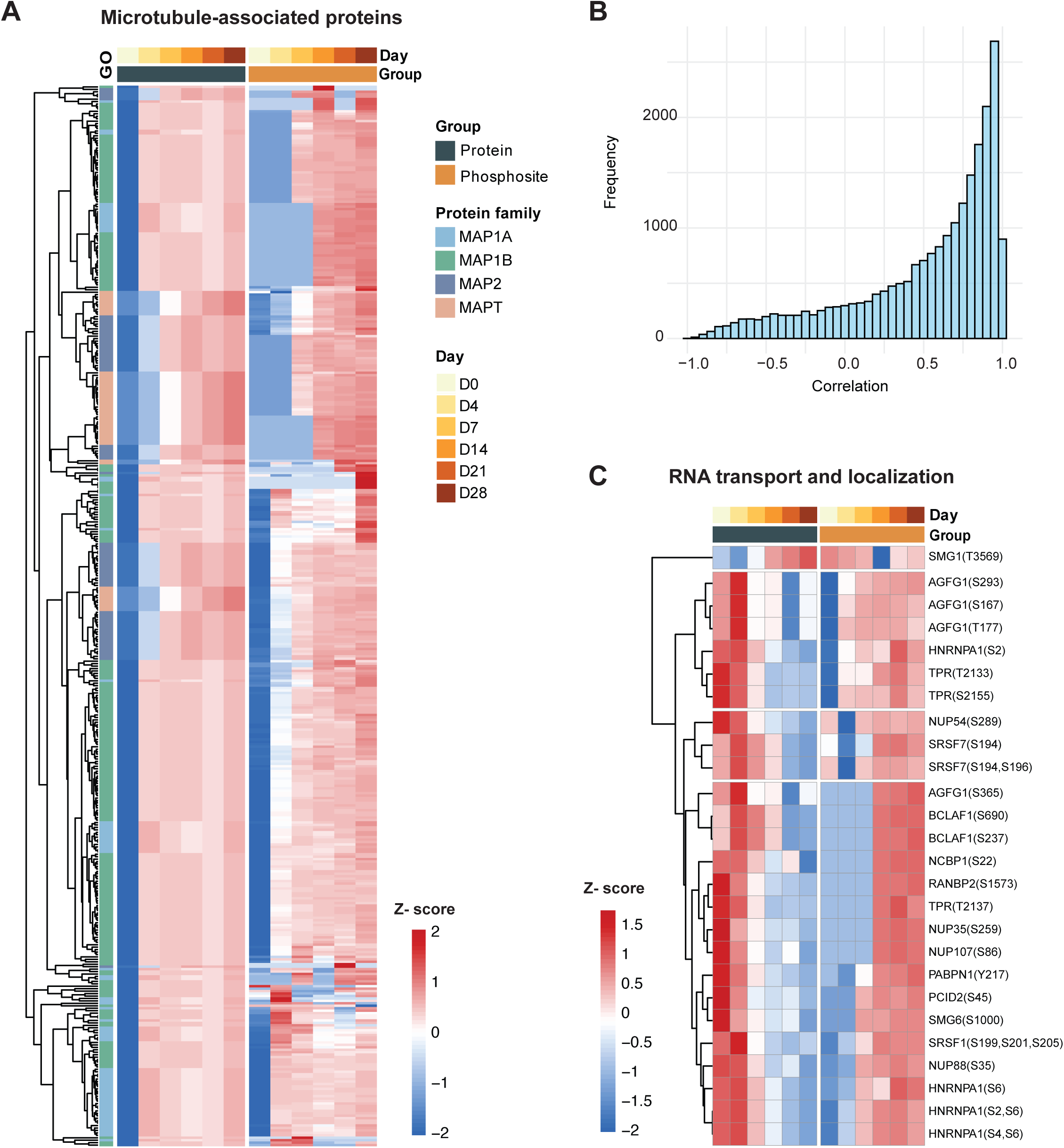
Dynamic trends in total protein and phosphosite levels within functional pathways. (A) Heatmap showing the dynamics of microtubule-associated proteins and their phosphosites. (B) Spearman correlation analysis between phosphosites and their corresponding unmodified proteins. (C) Discordant trends between the two omics data involve the RNA transport and localization pathway.

**Figure S4:**
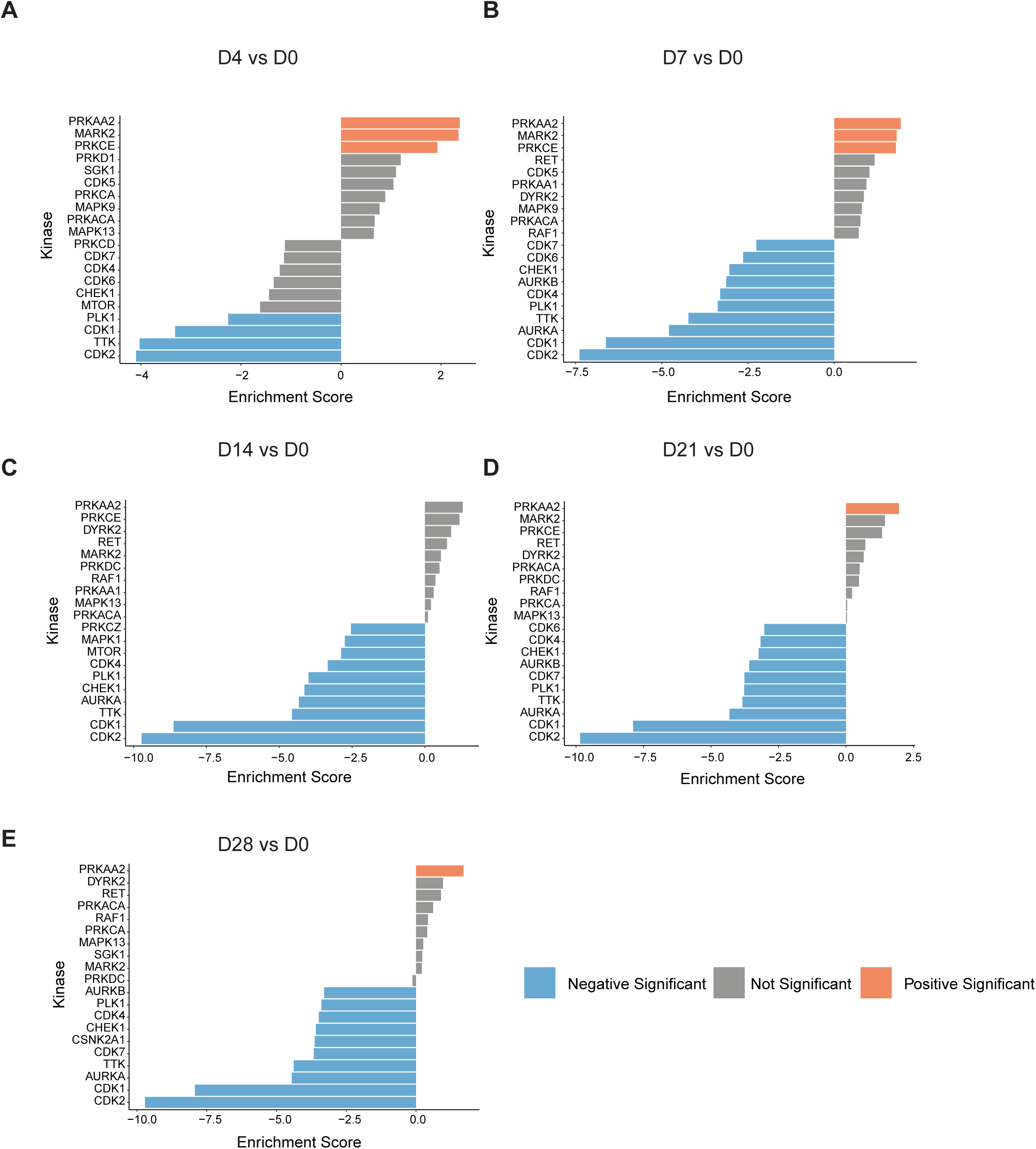
Time-point comparisons of kinase activity based on KSEA. (A) Bar plot showing the enrichment scores based on KSEA analysis comparing day D4 vs D0 (A) Bar plot showing the enrichment scores based on KSEA analysis comparing day D7 vs D0 (A) Bar plot showing the enrichment scores based on KSEA analysis comparing day D14 vs D0 (A) Bar plot showing the enrichment scores based on KSEA analysis comparing day D21 vs D0 (A) Bar plot showing the enrichment scores based on KSEA analysis comparing day D28 vs D0 Positive enrichment scores (orange bars) indicate kinases with increased activity, negative enrichment scores (blue bars) indicate kinases with decreased activity.

**Table S1 Protein abundance in the 513 proteomics data.**

**Table S2 Phosphopeptide abundance in the phosphoproteomics data.**

**Table S3 Kinase substrate enrichment score files**

